# Self-assembling SARS-CoV-2 nanoparticle vaccines targeting the S protein induces protective immunity in mice

**DOI:** 10.1101/2021.02.05.428685

**Authors:** Xingjian Liu, Haozhi Song, Jianmin Jiang, Xintao Gao, Yongzhu Yi, Yuting Shang, Jialei Li, Dan Li, Zhen Zeng, Yinü Li, Zhifang Zhang

## Abstract

The spike (S), a homotrimer glycoprotein, is the most important antigen target in the research and development of SARS-CoV-2 vaccine. There is no doubt that fully simulating the advanced structure of this homotrimer in the subunit vaccine development strategy is the most likely way to improve the immune protective effect of the vaccine. In this study, the preparation strategies of S protein receptor-binding domain (RBD) trimer, S1 region trimer, and ectodomain (ECD) trimer nanoparticles were designed based on ferritin nanoparticle self-assembly technology. The Bombyx mori baculovirus expression system was used to prepare these three nanoparticle vaccines with high expression levels in the silkworm. The immune results of mice show that the nanoparticle vaccine prepared by this strategy can not only induce an immune response by subcutaneous administration but also effective by oral administration. Given the stability of these ferritin-based nanoparticles vaccine, easy-to-use and low-cost oral immunization strategy can make up for the vaccination blind areas caused by the shortage of ultralow-temperature equipment and medical resources in underdeveloped areas. And the oral vaccine is also a very potential candidate to cut off the spread of SARS-CoV-2 in domestic and farmed animals, especially in stray and wild animals.

## Introduction

The coronavirus disease 2019 (COVID-19) has swept the globe for months. According to the World Health Organization (WHO) report, there are more than 400 million confirmed cases and more than5 million deaths as of 28 th February 2022(https://covid19.who.int/). The pathogen of COVID-19 was identified and named as Severe Acute Respiratory Syndrome Coronavirus 2 (SARS-CoV-2)^1^, which is another highly pathogenic human coronavirus that appeared after SARS-CoV (2003) and MERS-CoV (2012).

To cope with the spreading of the SARS-CoV-2 and COVID-19 Pandemic, deployment of effective physical protective measures and investment of large amounts of medical and health resources are indispensable. At the same time, developing an efficacious vaccine is also of paramount importance. The WHO maintains a document that includes most of the SARS-CoV-2 vaccines in development (https://www.who.int/publications/m/item/draft-landscape-of-covid-19-candidate-vaccines). According to this file, more than 300 vaccine candidates are currently in development.

Though preclinical studies of vaccines against SARS-CoV and MERS-CoV, the spike (S) protein, which is responsible for receptor binding and membrane fusion, has been identified to be the major antigenic target for coronavirus vaccines^2-4^. Researches about SARS-CoV-2 vaccines also confirm this point. Hence, most current vaccine candidates against SARS-CoV-2 target full-length or partial S protein^5^. Similar to HIV gp160 and influenza hemagglutinin, the S glycoprotein is a class I viral fusion protein that exists as a homotrimer with multiple glycosylation sites. Its immunogenicity was highly dependent on the advanced structure, which with great exposure to antigen epitopes^6-8^. Studies on the S-based subunit vaccines demonstrate that polymeric protein vaccines (dimer or trimer) were able to induce more neutralizing antibodies than monomer vaccines^9,10^. Although early vaccine development strategies involving subunit vaccines, especially trimeric subunit vaccines, have been less studied. However, with the investment of more scientific tools and scientific research forces, more and more studies have reported the research strategy of trimeric subunit vaccines^11-13^. In previous studies, Kanekiyo, M. constructed a self-assembling synthetic nanoparticle vaccine with influenza hemagglutinin trimer, which improves the potency and breadth of influenza virus immunity with lower immune dose^14^. This strategy provides an important reference for the development of S protein trimer vaccines.

In the present study, combining with the self-assembly characteristics of *Helicobacter pylori* ferritin, we have developed three SARS-CoV-2 nanoparticle vaccines, which containing receptor-binding domain (RBD) trimer, S1 trimer, and ectodomain (ECD) trimer, respectively. The BmNPV baculovirus expression system, reBmBac was used for the production^15^. The nanoparticle vaccines, prepared in silkworm, could induce protective immunity in mice not only by subcutaneous administration but also oral administration. It indicated that depend on the stability of ferritin nanoparticles, the nano-vaccines could be used to develop effective vaccination programs implemented in underdeveloped areas. Furthermore, in reference to the epidemic prevention strategy of rabies and influenza, animal vaccines against SARS-CoV-2 could also be developed using this method. Breaking transmission chains in animals, especially pets and farm animals is imperative^16^. By detecting the specific antibody titer and neutralizing antibody induced by these three nano-vaccines, the effectiveness of them was initially confirmed. Of which, the ECD-trimer nano-vaccine, which induced the most significant level of neutralizing antibody, was suitable for further development as a candidate vaccine.

## Results

### 1. Preparation of Self-assembling nanoparticle vaccines

The S glycoprotein nano-vaccines expression recombinant baculoviruses were prepared by using the reBmBac baculovirus expression system (BES). The RBD, S1 and ECD sequences of S protein were fusion expressed with *Helicobacter pylori* ferritin at N-terminal respectively (Fig. 1A). The fusion subunits produced in silkworms should self-assemble into 24-mer nanoparticles. And the S trimers were displayed on the nanoparticle surface (Fig. 1B). Western Blot analysis showed that the fusion proteins were successfully expressed (Fig. 1C). And the expression level of the fusion proteins, RBD-Fe, S1-Fe and ECD-Fe, were 187, 468 and 287 ug/mL larval haemolymph, respectively. Transmission electron microscopy (TEM) and Immunoelectron microscopy (IEM) confirmed the self-assembling of the three versions of nanoparticles: ECD-Nano, S1-Nano and RBD-Nano.

**Figure 1.**
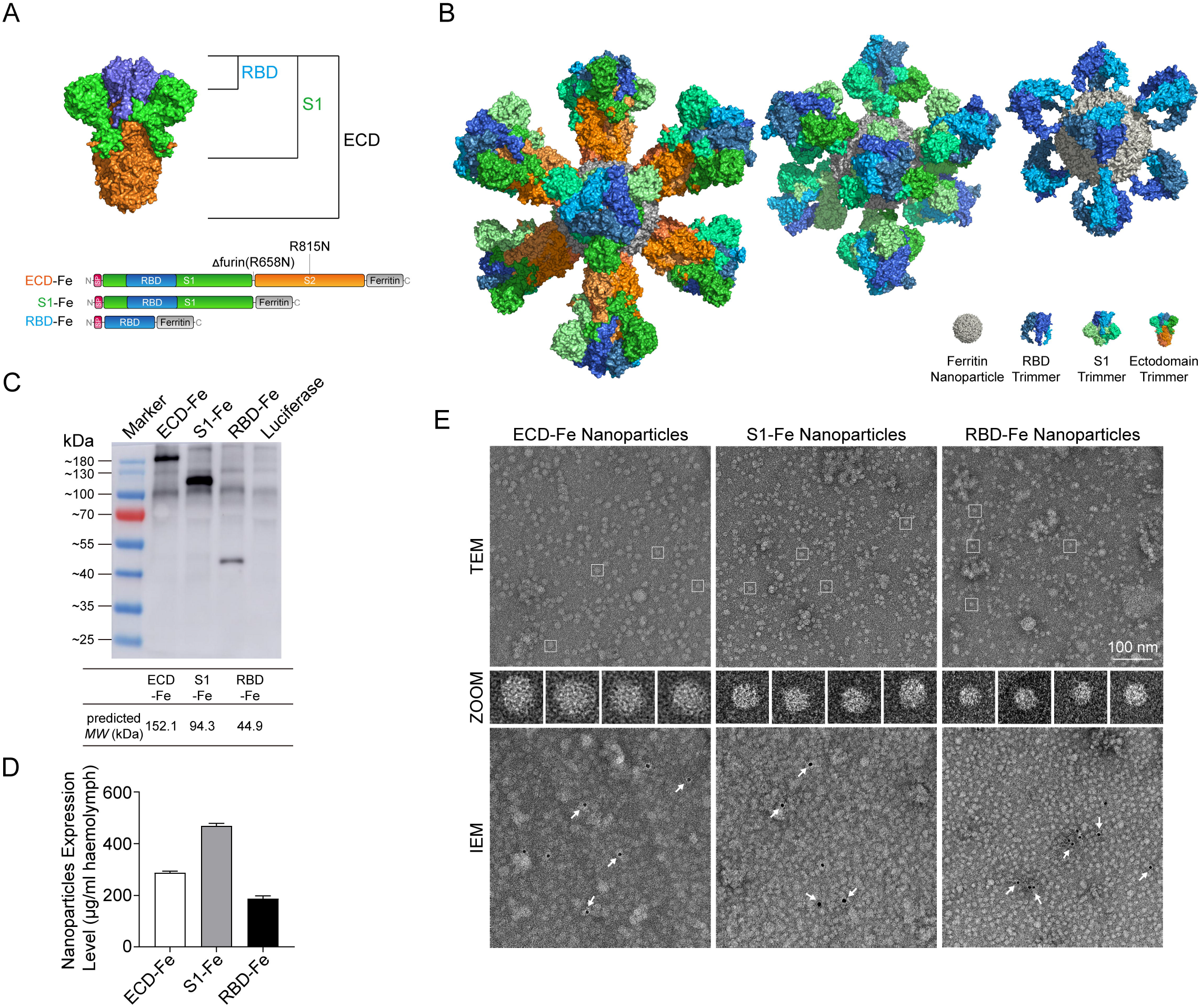
Construction and expression of nanoparticle vaccines. **(A)** Schematic of vaccine components which were RBD-Fe, S1-Fe and ECD-Fe. Fe: Ferritin, RBD: receptor-binding domain, ECD: ectodomain. **(B)** Schematic of nanoparticle of the three vaccines. PyMOL software was used for visualization^17^. **(C)** Western Blot analysis of the products prepared in silkworm by using an anti-RBD antibody. **(D)** ELISA of the S proteins to evaluate the expression levels. ECD-Fe with 287 ug/mL larval haemolymph, S1-Fe with 468 ug/mL larval haemolymph, RBD-Fe with 187 ug/mL larval haemolymph. **(E)** TEM image and two-dimensional (2D) reconstruction confirmed the successful assembly of each nanoparticle, IEM confirmed the S protein on the surface of the nanoparticle.

### 2. Immunogenicity of ECD-Nano, S1-Nano and RBD-Nano in mice

To evaluate the immunogenicity of these three nanoparticles, BALB/c mice were immunized twice at an interval of 3 weeks. Mice in subcutaneous administration groups (SA) were immunized with 10 μg of ECD-Nano or equal moles of S1-Nano and RBD-Nano. In proportion, mice in oral administration groups (OA) were immunized with 50 μg of the ECD-Nano or equal moles of S1-Nano and RBD-Nano. The sera were collected at 14 days after prime and boost. The titers of ECD-specific IgG, S1-specific IgG and RBD-specific IgG in sera were detected with ELISA using the corresponding recombinant protein of ECD, S1 or RBD, which produced and purified in *E. coli*. We found that the geometric mean titers (GMTs) of specific IgG elicited by ECD nanoparticle vaccines are 1,132 and 1,006 via subcutaneous and oral routes respectively 14 days after prime. And the GMTs of ECD specific IgG reach up to 8,072 and 4,528 35 days after prime. The GMTs of S1 specific IgG in S1-Nano immunized groups are 1,425 and 1,270 via subcutaneous and oral routes respectively at 14 days, and 9,057 and 5,701 at 35 days. The GMTs of RBD specific IgG in the other two groups are 1,794 and 1,006 via subcutaneous and oral routes respectively after prime, and 9,057 and 4,528 after boost. The antibody titers in OA groups are slightly lower than that in SA groups, but the best samples basically could reach 10^4^.

### 3. Nanoparticle vaccine elicited effective antibody responses in mice

To assess whether the sera contains effective neutralizing antibodies against SARS-CoV-2 infection, we firstly performed a confocal analysis to detect whether the specific antisera could block the interaction between nanoparticle and human angiotensin□converting enzyme 2 (hACE2). The 293T-hACE2 cells were hACE2-EGFP fusion protein overexpressed HEK293T cells. It is confirmed that the recombinant hACE2 was in the membrane of cells by colocalization analysis with EGFP (Fig. S1). As the confocal analysis results are shown in Fig. 3, antibodies in the antisera from mice immunized with different nano-vaccines could significantly inhibit the binding of RBD to hACE2 (Fig. 3).

**Figure 2.**
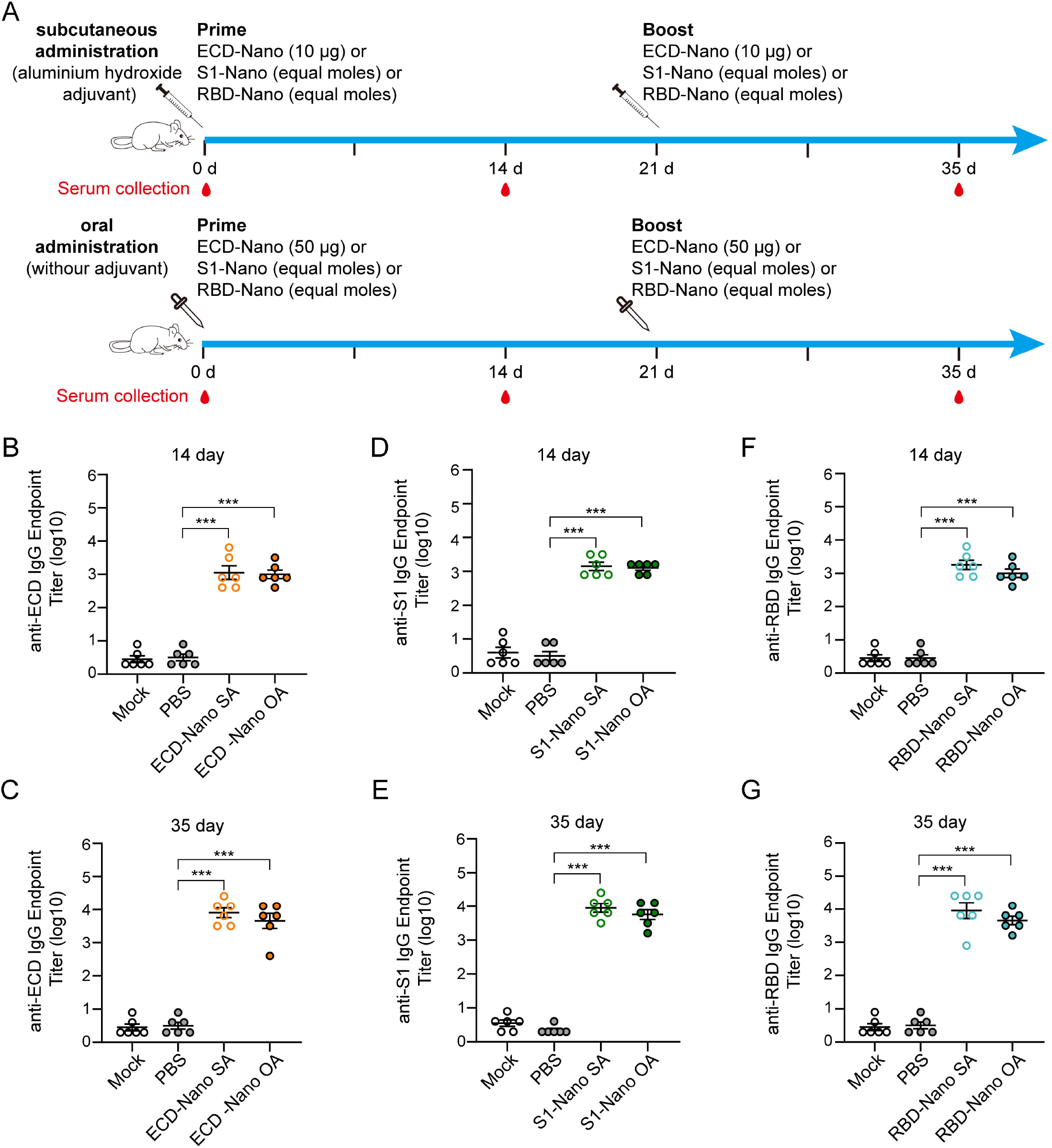
BALB/c mice immunized with nanoparticle vaccines produced SARS-CoV-2 ECD-S1- and RBD-specific antibodies. **(A)** The immunization protocols. Mice were immunized with vaccine via subcutaneous or oral administration. **(B)** and **(C)** Detection of ECD-specific IgG in different routes of administration of sera from day 14 and 35. **(D)** and **(E)** Detection of S1-specific IgG in different routes of administration of sera from day 14 and 35. **(F)** and **(G)** Detection of RBD-specific IgG in different routes of administration of sera from day 14 and 35.

**Figure 3.**
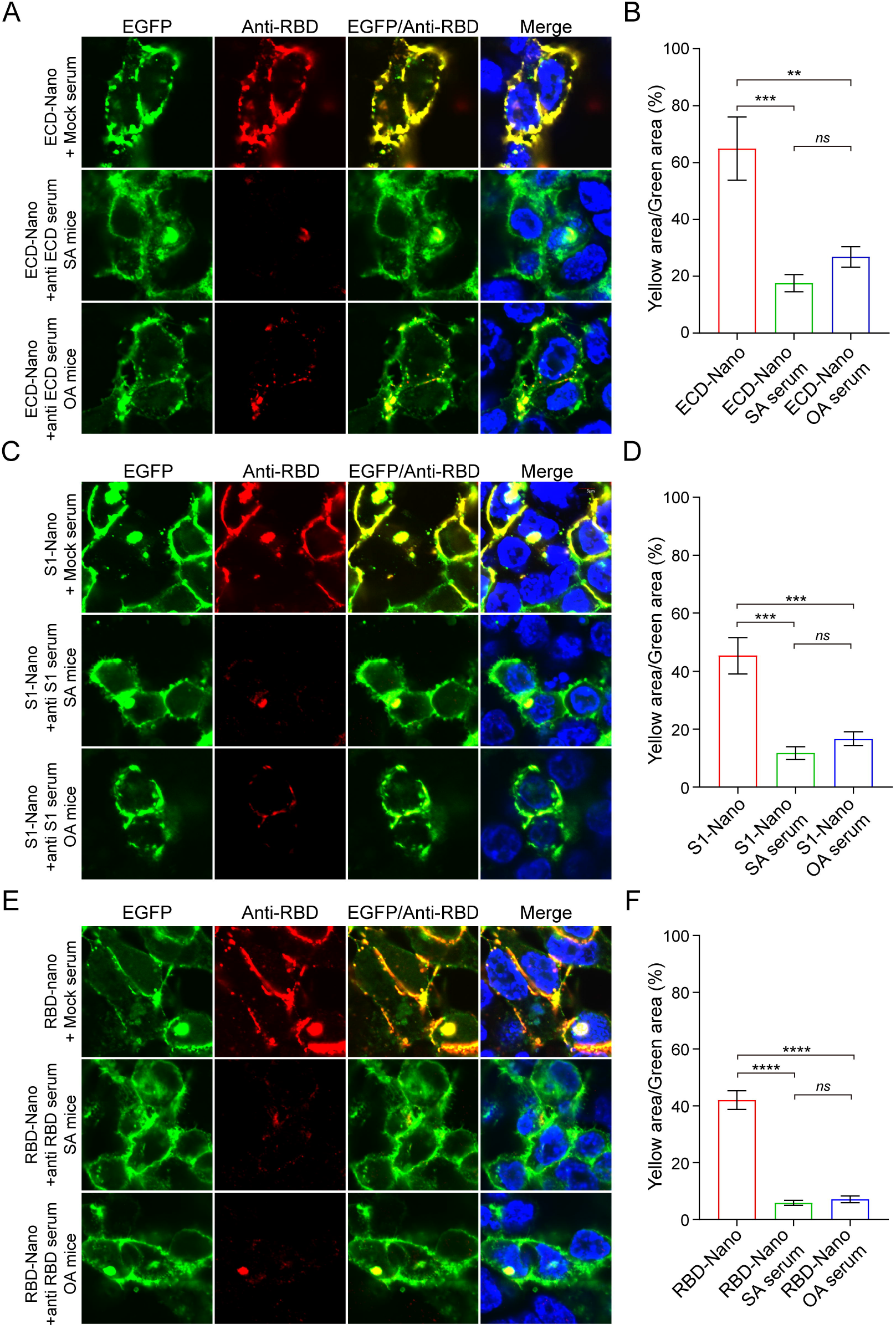
Inhibition of the binding of the S protein to hACE2. The ECD-Nano **(A)**, S1-Nano **(B)**, RBD-Nano **(C)** were added to 293T-hACE2 cells in the presence or absence of antisera at a dilution of 1:10. Confocal fluorescence analysis confirmed that all the three nanoparticles could specifically bind to hACE2 due to the existence of RBD structure. After treatment of the corresponding antiserum, the co-localization signal was significantly weakened, which indicated that the antibodies induced by mice immunized with nanoparticles vaccine had a certain neutralizing effect. ** p<0.01, *** p<0.001, **** p<0.0001.

To further investigate. We also tested the neutralization activity against authentic SARS-CoV-2 infection in cells. Origin stain and delta variants were both used. The sera were diluted by 2 times gradient dilution from 1:2. For origin stain, only the sera collection from ECD-Nano SA mice showed effective activity in VERO cells with a geometric mean 50% neutralization titers (NT50) of 160. For the delta variants, the neutralizing titer of ECD-Nano-induced antibody was 1:48, and the neutralization titer of S1-Nano-induced antibody was 1:32. However, due to the stringent operating conditions of the authentic virus, this neutralization test was only a preliminary verification. In the follow-up, it is still necessary to coordinate the experimental conditions for systematic comparative evaluation to determine whether each nanoparticle vaccines can induce the production of neutralizing antibodies via subcutaneous and oral immunization.

### 4. The oral route of this nanoparticle vaccine has a high level of tolerability and safety in mice

To explore the safety of oral immunization of nanoparticles vaccine, high-dose immunization experiments in mice were carried out. The total amount of immunity was increased to 5 times and was continuously immunized for 5 days. After 21 days of observation, the mice were euthanized, and the serum titer and tissue sections were determined. With a high dose of immunization, the body temperature and behavior of mice were normal, and no pathological characterization could be observed. The GMTs of ECD specific IgG in ECD-Nano immunized mice, S1 specific IgG in S1-Nano immunized mice and RBD specific IgG in RBD-Nano immunized mice are 2.8×10^4^, 2.8×10^4^, and 2.3×10^4^ at day 21 (Fig. S2). There were no obvious histopathologic changes in the organs of the three groups (Fig. S3).

## Discussion

In this study, we designed three versions of SARS-CoV-2 nanoparticle vaccines with different lengths of Spike respectively and evaluated the efficacy of them via two different administration routes of subcutaneous and oral. All three nanoparticle vaccines induced robust antibody responses against corresponding antigens. All of the nAbs could inhibit the binding of nanoparticles to hACE2. And the nAbs collected in ECD-Nano SA mice could inhibit authentic SARS-CoV-2 origin strain infection in cells with a nAb titer of 1:160. The nAbs of ECD-Nano- and S1-Nano-induced could inhibit SARS-CoV-2 delta varians infection in cells with a titer of 1:48 and 1:32 respectively. This result preliminarily showed that the SARS-CoV-2 nanoparticle vaccine constructed based on this strategy has a certain broad spectrum of immune protection against different strains. Of course, although the mice immunized orally in this study produced high specific antibodies, the neutralization effect of the virus in vitro experiments was not ideal. This may be related to the reasons that oral administration mainly causes mucosal immunity, and the related mechanism still needs further in-depth study.

Through preclinical studies on several coronaviruses, the spike protein has been identified as the best antigenic target for the development of vaccine^2,3^. SARS-CoV-2 Spike protein is also the best target for vaccine candidates^4,5^. Furthermore, it is generally believed that compared with monomers, subunit vaccines with native-structure-like trimer or at least homodimer of Spike protein were able to induce more neutralizing antibodies^10,18^. In the development strategy of ferritin-based nanoparticle vaccine, Spike subunits distributed evenly on the nanoparticle surface as native-like trimer structure. The characteristics of ferritin self-assembling nanoparticle also make up for the less of antigen density of Spike subunit^18,19^. It could improve the compatibility of S subunit with the antigen capture and presentation strategy of the host immune system^19,20^. Studies showed that the ferritin-based nanoparticle vaccines could induce high efficacy, broad-spectrum and persistent protection^14,21^. Two methods of linking Spike antigen to ferritin have been reported. One is to connect Spike to the surface of self-assembled ferritin nanoparticles by SpyTag-SpyCatcher (ST/SC) strategy^22^. The other is the fusion expression of Spike antigen and ferritin monomer for self-assembly^12^. Rather than the postprocessing linking strategy as reported such as ST/SC conjugation, we adopted the fusion expression strategy in which the S subunit was linked to the N-terminal of ferritin. By taking full advantage of BES on high-molecular-weight protein expression, this approach simplified the preparation of nanoparticles vaccine and improves the stability of nanoparticles. Besides, the ECD sequence used in this study can still encode stable S protein without introducing the two stabilizing proline mutations^23,24^. This suggested that the ferritin-based nanoparticles may improve the stability of recombinant S protein, which needs further study to confirm.

Gastrointestinal infection of SARS-CoV-2 has been confirmed^25,26^, which indicated Spike protein has resistance to the environment of the gastrointestinal tract. Additionally, ferritin nanoparticle displays remarkable thermal and chemical stability^27,28^. Based on these, we designed the oral administration group to investigate whether the nanoparticle vaccines could induce efficient antibody in mice. And the results showed that the oral route is feasible. We believe that this nanoparticle vaccine could be used as a competitive candidate for COVID-19 epidemic control in regions and countries with underdeveloped medical conditions. For a long time, Africa has long been a vulnerable area on prevention and control of infectious diseases^29^. According to statistical data, Africa has more than 2 million confirmed cases till the end of 2020. How to ensure that people in Africa and other low-income countries and regions enjoy the same vaccine protection is an important issue facing the world. Even though the approved vaccines have sufficient production capacity, low-temperature transportation and clean vaccination in these areas are still difficult to solve quickly and economically. The vaccines prepared based on the ferritin nanoparticle strategy in this study can be used as one of the candidates to solve this problem because of its stability and oral effectiveness.

On the other hand, oral immunization has more advantages in animal immunity, especially in stray animals or wild animals. Studies have confirmed that SARS-CoV-2 can infect mammals such as non-human primates, cats, dogs, ferrets, minks and hamsters^30-33^. And the virus is transmissible between cats and ferrets^30,34^. It has been reported that there are widespread infections of SARS-CoV-2 in the farmed mink population and showed a trend of reverse transmission to the community^16,35^. As a result, some countries have even decided to cull all the farmed minks^36,37^. WHO also pointed that mink appear to be “good reservoirs” for the disease, with a mutated strain having caused infections in a dozen people. It can be seen that the transmission of SARS-CoV-2 among animals is a great threat to farmers and the environment. Although several vaccines have been approved all over the world successively, it seems to be difficult to eliminate and cut off the infection and transmission of SARS-CoV-2 in animals in the short term, which is great resistance to the process of global epidemic prevention and control.

The nanoparticle vaccine in this study has low production cost and has been proved to be able to induce the body to produce high titers of specific antibodies by oral route. It can be used not only as a human oral vaccine similar to OPV and as an oral candidate vaccine, but also as an effective scheme for large-scale immunization of pets and farmed animals, especially stray animals or wild animals, to eliminate and cut off the intermediate source of infection and avoid the mutation caused by the long-term transmission of the virus in animals, which is of great significance to the prevention and control of the global epidemic and the promotion of immunization programs. In this way, the intermediate source of infection of animals can be eliminated and cut off, and the mutation caused by the long-term spread of the virus in animals can also be avoided. This is of great significance to the prevention and control of the global epidemic and the promotion of the immunization process. Baculovirus expression system (BES) is suitable for the expression of eukaryotic-derived proteins. Many BES-based SARS-CoV-2 vaccine development researches have confirmed that the BES is also very suitable for the expression of S protein^9^. The silkworm BES used in this study can be used to prepare nanoparticles vaccine in silkworm, but not in cells. It has the advantages of low production cost and high expression. It is generally believed that the expression of 5 silkworm pupae is equivalent to that of 1 liter of insect cells^38^. The recombinant baculovirus strains used in this study also have the potential to further increase the expression level by 2-5 times through a series of optimization measures^15,39^. The advantages of low cost and high expression of BmNPV BES and the effectiveness of nanoparticles vaccine orally provide a sufficient guarantee for high-dose oral immunization to control the spread of SARS-CoV-2 in animals.

The results of the high-dose immune experiment preliminarily verified the safety of oral immunization of the nanoparticles vaccine developed in this study. High-dose continuous immunization induced mice to produce efficient specific immunity, and the antibody titer was even higher than that of the injection group. This suggesting that the subjects have a high level of tolerability to the immune dose and frequency of oral route in practical application. This provides ample optimization space for exploring the optimization of the oral vaccine immunization program.

In this study, we found that in the preliminary virus neutralization experiment in vitro, ECD-Nano induced antibodies showed a significant neutralization effect. This may be due to the fact that ECD contains both S1 and S2 extracellular regions, which can induce a richer variety of neutralizing antibodies. It is more effective to inhibit the virus from entering the cell. The structure of S protein of SARS-CoV-2 is similar to that of SARS-CoV and MERS-CoV. The RBD of S1 is responsible for receptor recognition and binds to ACE2, which is the preferred antigen for many subunit vaccine development strategies. The HR region of S2 is responsible for membrane fusion mediating the entry of viruses into cells, and the sequence is relatively conservative, which is considered to be a promising target for the development of fusion inhibitors against SARS-CoV-2 infection^40^. However, the S protein of SARS-CoV-2 also has its particularity. Its RBD conformation changes dynamically between exposed and hidden states, which makes it easier to evade immune surveillance against RBD^6,7^. There is a furin cleavage site between S1/S2, which can mediate the pre-activation caused by proprotein converting enzyme, making it easier for the virus to enter the host cell. That is to say, SARS-CoV-2 can also enter the host cell when S2 is directly exposed^6^. Some studies also have shown that antibodies against RBD or HR region alone can inhibit the entry of SARS-CoV-2 into cells^18^. Thus, vaccines containing both RBD and HR will theoretically have better immune protection. The ECD-Nano and S1-Nano vaccine in this study contains more antigenic epitopes than those of RBD-Nano. And its structure is more similar to the natural trimer of S protein. So, it may be more suitable as a candidate vaccine development strategy.

## Materials and Methods

### Reagents

Rabbit anti-SARS-CoV-2 spike ECD polyclonal IgG, Rabbit anti-SARS-CoV-2 spike S1 polyclonal IgG, Rabbit anti-SARS-CoV-2 spike RBD polyclonal IgG was purchased from Sino Biological (Beijing, CN). HRP-conjugated goat anti-rabbit IgG was purchased from ZSGB-BIO (Beijing, CN). CF568 labeled Goat anti-rabbit IgG (H+L) was purchased from Biotium Inc. (Hayward, CA, USA). The luciferase assay system was obtained from Promega (Madison, WI, USA). Goat Anti-Rabbit IgG (whole molecule) Gold antibody was obtained from Sigma Aldrich (St. Louis, MO, USA).

### Plasmids, bacterial strains and cell lines

The defective-rescue BmNPV BES reBmBac vector was constructed and maintained in our laboratory^15^. The pET-28a (+) vector was saved in our laboratory. The recombinant BmNPV transfer vector pBmPH (GenBank submission ID: 2426173) was constructed and maintained in our laboratory. BmN cells was preserved in our laboratory and cultured in TC100 insect cell culture medium (Applichem, Darmstadt, Germany) with 10% fetal bovine serum (FBS, Gibco, USA) at 27°C. HEK-293T cells was obtained from ATCC and cultured in DMEM (Thermo Fisher, Waltham, MA, USA) with 10% FBS at 37°C. The *Escherichia coli* strains Top10 and BL21 (DE3) were cultured in Luria-Bertani (LB) medium.

### Recombinant BmNPV construction

Three peptide sequences of nanoparticle vaccines were designed according to the full-length sequence of S protein (RefSeq: YP_009724390.1). The sequences of ECD (14-1213 aa, R685N), S1 region (14-685 aa, R685N) and RBD (319-541 aa) were linked with *Helicobacter pylori* ferritin (RefSeq: WP_000949190.1, 5-167 aa, N19Q) via SGG linker. The human T-cell surface glycoprotein CD5 signal peptide (RefSeq: NP_055022.2, 1-24 aa) was used as signal peptide of these three sequences. After codon preference optimization, the three designed sequences were synthesized by GenScript (Nanjing, CN) with *Bam*HI and *Eco*RI restriction sites at the upstream and downstream, respectively. The ECD-Ferritin sequence was inserted into pBmPH vector through *Bam*HI/*Eco*RI digestion to construct the transfer vector pBmPH-ECD-Fe. The pBmPH-S1-Fe and pBmPH-RBD-Fe vectors were constructed the same way. The transfer vectors were then cotransfected in BmN cells with reBmBac respectively. The recombinant BmNPV baculovirus, reBm-ECD-Fe, reBm-S1-Fe and reBm-RBD-Fe, were obtained in the cell supernatant of cotransfection.

### Nanoparticles expression and purification

Fifth instar silkworm larvae or pupae were injected with recombinant viruses (5 μL of the supernate, approximately 0.5×10^5^ pfu) between the abdominal knobs on the backside and then reared or incubated at 25-27°C and 65% humidity. Larval haemolymph or pupae body expressing nanoparticles were harvested after 108-120 h. The crude samples were purified by gentle ultrasonic crushing and ultracentrifugation by 30% sucrose cushion at 120,000×g for 3 h. The white transparent nanoparticle pellet was resuspended in PBS overnight. The concentrations of the nanoparticles were detected by ELISA using standard samples of ECD, S1 and RBD (Sino Biological, Beijing, CN). Whether the nanoparticles were assembled successfully or not was observed by transmission electron microscope (TEM). Immunoelectron microscopy (IEM) was used to identify whether there was S protein on the surface of nanoparticles.

### Animal ethics statement

The mouse immunogenicity studies were performed in Biotechnology Research Institute and Institute of Animal Sciences of Chinese Academy of Agricultural Science. The Ethics Review Board of Biotechnology Research Institute and Institute of Animal Sciences approved this study. All animal procedures were in strict accordance with the guidelines of the Animal Care and Use Committee of Biotechnology Research Institute and Institute of Animal Sciences.

### Mouse study designs

Six BALB/c mice (6-8 weeks of age) were assigned in every group. Mice in Subcutaneous Administration (SA) group were subcutaneously immunized with 10μg dose of crude purified nanoparticle vaccines ECD-Nano, or equal moles dose of S1-Nano or RBD-Nano, formulated with aluminium hydroxide adjuvant. Mice in Oral Administration (OA) group were fed immunized with 50 μg dose of crude expression products ECD-Nano, or equal moles dose of S1-Nano or RBD-Nano, without adjuvant. All the mice were vaccinated with the above vaccines in a prime/boost manner which was vaccinating mice on days 0 and day 21. Sera were collected twice on days 14 and 35. Mice were euthanized at week 6. PBS administrator Mice were the control group.

In safety test groups, mice were fed with 50 μg dose of nanoparticles per day for five days continuously. Sera were collected on days 0 and 21. Mice were euthanized and dissected on day 21.

### Identification of serum antibodies against the Spike in mice using an ELISA assay

Blood samples were collected from the retro-orbital plexus of mice on days 14 and 35. After coagulation at 37^°^C for 2 h, blood samples were centrifuged at 2,000 rpm for 10 min. The serum was collected and stored at -20^°^C. Recombinant ECD-His, S1-His, and RBD-His expressed and purified from *E*.*coli* BL21 (DE3), were used to coat flat-bottom 96-well plates at a final concentration of 1 μg/mL. After overnight incubation at 4^°^C, plates were washed three times with PBST and added with blocking buffer containing 1% BSA in PBST, followed by 1 h incubation at room temperature. Mouse sera were serially diluted and added to the plates. After 1 h incubation at 37^°^C, the plates were washed three times and added the HRP-conjugated Goat anti-mouse IgG antibody with a dilution of 1:5000. After incubation for 1 h at 37^°^C, the plates were washed three times with PBST and developed with TMB (Solarbio, Beijing, CN) for 10 min. The reactions were stopped with a stop solution (1M H_2_SO_4_). The absorbance at 450 nm was measured on a microplate reader.

### Measurement of the inhibition of RBD binding to cell-surface ACE2

The hACE2 was co-expressed with EGFP using pEGFP-N3 vector in HEK-293T cells. The purified Spike nanoparticles were incubated with PBS or antisera for 1 h respectively. The mixtures were then added into the 293T-hACE2 cells, followed by incubating for 12 h. The cells were fixed with paraformaldehyde buffer, followed by incubating with Rabbit anti-SARS-CoV-2 spike RBD polyclonal IgG for 2 h at 37^°^C. After washing, the cells were incubated with CF568 labeled Goat anti-rabbit IgG (H+L) with a dilution of 1:5000 for 2 h at 37^°^C. Images in fluorescence were acquired by confocal scanning microscopy.

### SARS-CoV-2 authentic virus neutralization assay

To assess the neutralization of the SARS-CoV-2 infection in vitro, Vero cells (5×10^4^) were seeded in 96-well plates and grown overnight. SARS-CoV-2 origin stain and delta variants (approximately 100TCID_50_) was preincubated with an equal volume of sera (serially diluted from 1:2) from immunized mice. After incubation at 37^°^C for 1h, the mixture was added to Vero cells. After 3 days, cytopathogenic effects of the cells were recorded under the microscope. The neutralization assay was carried out in BSL-3.

### Statistical analysis

All statistical analyses were carried out with GraphPad Prism software. P□<□0.05 was considered significant. *P□<□0.05, **P□<□0.01, ***P□<□0.001, ****P□<□0.0001.

## Acknowledgments

This work was supported by National Natural Sciences Foundation of China (No. 32072796 and 32002236), the National Transgenic Major Project of China (2019ZX08010-004) and The Agricultural Science and Technology Innovation Program.

## Ethics statement

None required

## Reference

1 Coronaviridae Study Group of the International Committee on Taxonomy of, V. The species Severe acute respiratory syndrome-related coronavirus: classifying 2019-nCoV and naming it SARS-CoV-2. Nat Microbiol 5, 536–544, doi:10.1038/s41564-020-0695-z (2020).

2 Graham, R. L., Donaldson, E. F. & Baric, R. S. A decade after SARS: strategies for controlling emerging coronaviruses. Nat Rev Microbiol 11, 836–848, doi:10.1038/nrmicro3143 (2013).

3 Yong, C. Y., Ong, H. K., Yeap, S. K., Ho, K. L. & Tan, W. S. Recent Advances in the Vaccine Development Against Middle East Respiratory Syndrome-Coronavirus. Front Microbiol 10, 1781, doi:10.3389/fmicb.2019.01781 (2019).

4 Krammer, F. SARS-CoV-2 vaccines in development. Nature 586, 516–527, doi:10.1038/s41586-020-2798-3 (2020).

5 Amanat, F. & Krammer, F. SARS-CoV-2 Vaccines: Status Report. Immunity 52, 583–589, doi:10.1016/j.immuni.2020.03.007 (2020).

6 Shang, J. et al. Cell entry mechanisms of SARS-CoV-2. Proc Natl Acad Sci U S A 117, 11727–11734, doi:10.1073/pnas.2003138117 (2020).

7 Fan, X., Cao, D., Kong, L. & Zhang, X. Cryo-EM analysis of the post-fusion structure of the SARS-CoV spike glycoprotein. Nat Commun 11, 3618, doi:10.1038/s41467-020-17371-6 (2020).

8 Liu, L. et al. Potent neutralizing antibodies against multiple epitopes on SARS-CoV-2 spike. Nature 584, 450–456, doi:10.1038/s41586-020-2571-7 (2020).

9 Tian, J. H. et al. SARS-CoV-2 spike glycoprotein vaccine candidate NVX-CoV2373 immunogenicity in baboons and protection in mice. Nat Commun 12, 372, doi:10.1038/s41467-020-20653-8 (2021).

10 Dai, L. et al. A Universal Design of Betacoronavirus Vaccines against COVID-19, MERS, and SARS. Cell 182, 722–733 e711, doi:10.1016/j.cell.2020.06.035 (2020).

11 Joyce, M. G. et al. SARS-CoV-2 ferritin nanoparticle vaccines elicit broad SARS coronavirus immunogenicity. Cell Rep 37, 110143, doi:10.1016/j.celrep.2021.110143 (2021).

12 Powell, A. E. et al. A Single Immunization with Spike-Functionalized Ferritin Vaccines Elicits Neutralizing Antibody Responses against SARS-CoV-2 in Mice. ACS Cent Sci 7, 183–199, doi:10.1021/acscentsci.0c01405 (2021).

13 Wang, W., Huang, B., Zhu, Y., Tan, W. & Zhu, M. Ferritin nanoparticle-based SARS-CoV-2 RBD vaccine induces a persistent antibody response and long-term memory in mice. Cell Mol Immunol 18, 749–751, doi:10.1038/s41423-021-00643-6 (2021).

14 Kanekiyo, M. et al. Self-assembling influenza nanoparticle vaccines elicit broadly neutralizing H1N1 antibodies. Nature 499, 102–106, doi:10.1038/nature12202 (2013).

15 Liu, X. et al. A Highly Efficient and Simple Construction Strategy for Producing Recombinant Baculovirus Bombyx mori Nucleopolyhedrovirus. PLoS One 11, e0152140, doi:10.1371/journal.pone.0152140 (2016).

16 Oude Munnink, B. B. et al. Transmission of SARS-CoV-2 on mink farms between humans and mink and back to humans. Science 371, 172–177, doi:10.1126/science.abe5901 (2021).

17 DeLano, W. L. Pymol: An open-source molecular graphics tool. CCP4 Newsletter on protein crystallography 40, 82–92.

18 Ma, X. et al. Nanoparticle Vaccines Based on the Receptor Binding Domain (RBD) and Heptad Repeat (HR) of SARS-CoV-2 Elicit Robust Protective Immune Responses. Immunity 53, 1315–1330 e1319, doi:10.1016/j.immuni.2020.11.015 (2020).

19 Wang, N., Shang, J., Jiang, S. & Du, L. Subunit Vaccines Against Emerging Pathogenic Human Coronaviruses. Front Microbiol 11, 298, doi:10.3389/fmicb.2020.00298 (2020).

20 Shin, M. D. et al. COVID-19 vaccine development and a potential nanomaterial path forward. Nat Nanotechnol 15, 646–655, doi:10.1038/s41565-020-0737-y (2020).

21 Georgiev, I. S. et al. Two-Component Ferritin Nanoparticles for Multimerization of Diverse Trimeric Antigens. ACS Infect Dis 4, 788–796, doi:10.1021/acsinfecdis.7b00192 (2018).

22 Kang, Y. F. et al. Rapid Development of SARS-CoV-2 Spike Protein Receptor-Binding Domain Self-Assembled Nanoparticle Vaccine Candidates. ACS Nano, doi:10.1021/acsnano.0c08379 (2021).

23 Wrapp, D. et al. Cryo-EM structure of the 2019-nCoV spike in the prefusion conformation. Science 367, 1260–1263, doi:10.1126/science.abb2507 (2020).

24 Pallesen, J. et al. Immunogenicity and structures of a rationally designed prefusion MERS-CoV spike antigen. Proc Natl Acad Sci U S A 114, E7348–E7357, doi:10.1073/pnas.1707304114 (2017).

25 Xiao, F. et al. Evidence for Gastrointestinal Infection of SARS-CoV-2. Gastroenterology 158, 1831–1833 e1833, doi:10.1053/j.gastro.2020.02.055 (2020).

26 Zang, R. et al. TMPRSS2 and TMPRSS4 promote SARS-CoV-2 infection of human small intestinal enterocytes. Sci Immunol 5, doi:10.1126/sciimmunol.abc3582 (2020).

27 He, D. & Marles-Wright, J. Ferritin family proteins and their use in bionanotechnology. N Biotechnol 32, 651–657, doi:10.1016/j.nbt.2014.12.006 (2015).

28 Wang, W. et al. Ferritin nanoparticle-based SpyTag/SpyCatcher-enabled click vaccine for tumor immunotherapy. Nanomedicine 16, 69–78, doi:10.1016/j.nano.2018.11.009 (2019).

29 Nkengasong, J. N., Ndembi, N., Tshangela, A. & Raji, T. COVID-19 vaccines: how to ensure Africa has access. Nature 586, 197–199, doi:10.1038/d41586-020-02774-8 (2020).

30 Halfmann, P. J. et al. Transmission of SARS-CoV-2 in Domestic Cats. New Engl J Med 383, 592–594, doi:10.1056/NEJMc2013400 (2020).

31 Bosco-Lauth, A. M. et al. Experimental infection of domestic dogs and cats with SARS-CoV-2: Pathogenesis, transmission, and response to reexposure in cats. Proc Natl Acad Sci U S A 117, 26382–26388, doi:10.1073/pnas.2013102117 (2020).

32 Schlottau, K. et al. SARS-CoV-2 in fruit bats, ferrets, pigs, and chickens: an experimental transmission study. Lancet Microbe 1, e218–e225, doi:10.1016/S2666-5247(20)30089-6 (2020).

33 Shi, J. et al. Susceptibility of ferrets, cats, dogs, and other domesticated animals to SARS-coronavirus 2. Science 368, 1016–1020, doi:10.1126/science.abb7015 (2020).

34 Richard, M. et al. SARS-CoV-2 is transmitted via contact and via the air between ferrets. Nat Commun 11, 3496, doi:10.1038/s41467-020-17367-2 (2020).

35 Munnink, B. B. O. et al. Jumping back and forth: anthropozoonotic and zoonotic transmission of SARS-CoV-2 on mink farms. bioRxiv (2020).

36 Koopmans, M. SARS-CoV-2 and the human-animal interface: outbreaks on mink farms. Lancet Infect Dis 21, 18–19, doi:10.1016/S1473-3099(20)30912-9 (2021).

37 Oreshkova, N. et al. SARS-CoV-2 infection in farmed minks, the Netherlands, April and May 2020. Euro Surveill 25, doi:10.2807/1560-7917.ES.2020.25.23.2001005 (2020).

38 Usami, A. et al. Comparison of recombinant protein expression in a baculovirus system in insect cells (Sf9) and silkworm. J. Biochem. 149, 219–227, doi:10.1093/jb/mvq138 (2011).

39 Liu, X. et al. A construction strategy for a baculovirus-silkworm multigene expression system and its application for coexpression of type I and type II interferons. Microbiologyopen 9, e979, doi:10.1002/mbo3.979 (2020).

40 Huang, Y., Yang, C., Xu, X. F., Xu, W. & Liu, S. W. Structural and functional properties of SARS-CoV-2 spike protein: potential antivirus drug development for COVID-19. Acta Pharmacol Sin 41, 1141–1149, doi:10.1038/s41401-020-0485-4 (2020).

