## Supplementary material for "Self-assembling SARS-CoV-2 nanoparticle vaccines targeting the S protein induces protective immunity in mice": (Fig. S1)

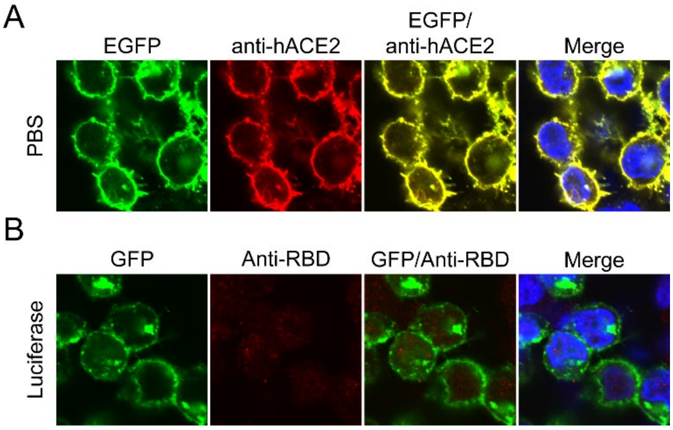


Figure S1. **(A)** The human ACE2 was co-expressed with EGFP in HEK-293T cells. **(B)** Baculovirus and other proteins in silkworm did not bind with hACE2.
