## Supplementary material for "Self-assembling SARS-CoV-2 nanoparticle vaccines targeting the S protein induces protective immunity in mice": (Fig. S2)

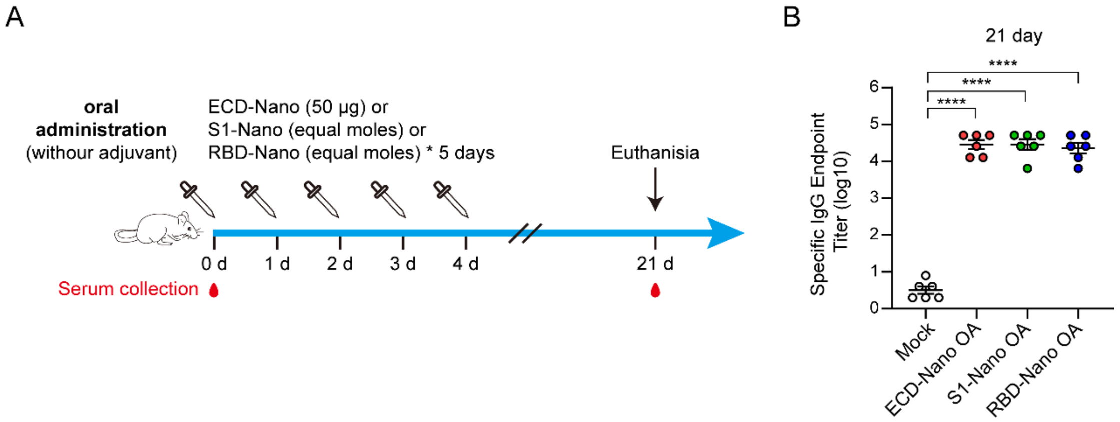


Figure S2. Safety assay of nanoparticle vaccines in mice via oral administration with high-dose. (A) The immunization protocols. (B) Detection of the specific IgG of sera of the oral immunized mice. The GMTs of ECD specific IgG in ECD-Nano immunized mice and S1 specific IgG in S1-Nano immunized mice were 28,707. The GMT of RBD specific IgG in RBD-Nano immunized mice was 22,803.
